# Integrin β3, a RACK1 interacting protein, is required for porcine reproductive and respiratory syndrome virus infection and NF-κB activation in Marc-145 cells

**DOI:** 10.1101/2020.01.27.922476

**Authors:** Chao Yang, Rui Lan, Xiaochun Wang, Qian Zhao, Xidan Li, Junlong Bi, Jing Wang, Guishu Yang, Yingbo Lin, Jianping Liu, Gefen Yin

**Author notes:** Although shared co-first author. Chao Yang performed more experiments than Rui Lan. These authors contributed equally to this work (Chao Yang, Rui Lan). Correspondence: Correspondence and requests for materials should be addressed to J.L. or G.Y.

## Abstract

Porcine reproductive and respiratory syndrome virus (PRRSV) is the pathogen of porcine reproductive and respiratory syndrome (PRRS), which is one of the most economically harmful diseases in modern pig production worldwide. Receptor of activated protein C kinase 1 (RACK1) was previously shown to be indispensable for the PRRSV replication and NF-κB activation in Marc-145 cells. Here we identified a membrane protein, integrin β3 (ITGB3), as a RACK1-interacting protein. PRRSV infection in Marc-145 cells upregulated the ITGB3 expression. Abrogation of ITGB3 by siRNA knockdown or antibody blocking inhibited PRRSV infection and NF-κB activation, while on the other hand, overexpression of ITGB3 enhanced PRRSV infection and NF-κB activation. Furthermore, inhibition of ITGB3 alleviated the cytopathic effects and reduced the TCID_50_ titer in Marc-145 cells. We also showed that RACK1 and ITGB3 were NF-κB target genes during PRRSV infection, and that they regulate each other. Our data indicate that ITGB3, presumably as a co-receptor, plays an imperative role for PRRSV infection and NF-κB activation in Marc-145 cells. PRRSV infection activates a positive feedback loop involving the activation of NF-κB and upregulation of ITGB3 and RACK1 in Marc-145 cells. The findings would advance our elaborated understanding of the molecular host–pathogen interaction mechanisms underlying PRRSV infection in swine and suggest ITGB3 and NF-κB signaling pathway as potential therapeutic targets for PRRS control.

**IMPORTANCE:** Porcine reproductive and respiratory syndrome virus (PRRSV) is one of the pathogens in pig production worldwide. Several cell surface receptors, such as heparan sulphate, sialoadhesin, vimentin and CD163, were identified to be involved in PRRSV infection in porcine alveolar macrophages (PAMs). We identified a cell surface protein, integrin β3 (ITGB3), as an interacting protein with receptor of activated protein C kinase 1 (RACK1) from Marc-145 cells. ITGB3 interacts with RACK1 and facilitates PRRSV infection and NF-κB activation in Marc-145 cells, presumably as a co-receptor of CD136 or vimentin. Both ITGB3 and RACK1 were NF-κB target genes, and they regulate each other. The activation of NF-κB and the transcription of its downstream genes are beneficial for PRRSV infection/replication. The novel findings would advance our elaborated understanding of the molecular host–pathogen interaction mechanisms underlying PRRSV infection in swine and suggest ITGB3-RACK1-NF-κB axis as a potential therapeutic target for PRRS control.

---

Porcine reproductive and respiratory syndrome (PRRS) is characterized by respiratory disorders in piglets and reproductive failure in sows (1). The contributing pathogen, porcine reproductive and respiratory syndrome virus (PRRSV), is a swine-specific enveloped single-stranded positive-sense RNA virus that belongs to the family *Arteriviridae*, genus *Arterivirus* (2). Since the first PRRS outbreak in the united states in 1987, PRRSV infections is nowadays detectable in almost all swine-producing countries, causing one of the highest economic impacts in modern pig production worldwide (3). Several cell surface receptors that are involved in PRRSV infection in porcine alveolar macrophages (PAMs) has been identified, including heparan sulphate (HS) (4), sialoadhesin (Sn) (5) and CD163 (6, 7).

Integrins are a family of heterodimer transmembrane receptors consisting of eighteen α-subunits and eight β-subunits that play pivotal roles in the binding of cells to extracellular matrix (ECM) (8, 9), transmembrane signal transduction (10, 11) and immune responses (12). Cellular integrins are common receptors exploited by diverse viral pathogens for cell entry and infection (13), such as Epstein-Barr virus (EBV)(14), human immunodeficiency virus 1 (HIV-1) (15, 16) and Simian virus 40 (17). The signalling functions of integrins are conducted through the formation of signalling complexes at the cytoplasmic face of the plasma membrane with its cofactors, for example focal adhesion kinase (FAK) (18) and receptor of activated protein C kinase 1 (RACK1) (19, 20).

RACK1, a 36-kDa protein comprising of seven WD-40 repeats, is well known as a scaffolder protein (21) and plays its unique functions in various cancers (22–24) and infections (24–28). RACK1 serves as an adaptor molecule for the binding of key signaling molecules (19, 22–24, 29–31),with elaborate involvement in the regulation of multiple signal pathways (24, 30, 32). RACK1 is reported to interact with β integrins to stabilize the focal adhesion through cell-ECM interaction with the entanglement of integrin-induced FAK autophosphoryation (19, 31, 33).

Nuclear factor kappa beta (NF-κB), whose activation by a variety of signals regulates the expression of many target genes, is a key regulator of cellular events. NF-κB signaling pathway modulates a broad range of biological processes including immune response, inflammation, cell proliferation, tumorigenesis and apoptosis. Binding of the viral particle to its receptors and the accumulation of viral products, including dsRNA and viral proteins, activate NF-κB signaling cascades through various processes. Multiple viruses, including HIV (34, 35), herpesviruses (36), and hepatitis C virus (HCV) (37), have in turn evolved sophisticated strategies to alert NF-κB signaling.

Our previous studies (26, 28) showed that RACK1 is indispensable for PRRSV replication and NF-κB activation in Marc-145 cells. As a build-up of that concept, the current study identified integrin β3 (ITGB3) as a interacting factor of RACK1 during the PRRSV infection in Marc-145 cells. Our data indicate that PRRSV infection upregulates the expression of RACK1 interacting ITGB3, which participates in the activation of the NF-κB signaling pathway, which in turn promotes PRRSV infection, presumably through regulation of RACK1 and ITGB3. The findings can advance our elaborated understanding of the molecular host–pathogen interaction mechanisms of PRRSV and suggest ITGB3 and NF-κB signaling pathway as potential therapeutic targets for PRRS control.

## RESULTS

### ITGB3 was identified as a RACK1 interacting protein

Venny diagram analysis (https://bioinfogp.cnb.csic.es/tools/venny/) of the four datasets from the pulldown-MS assay (Supplementary Table 1) revealed that there were 179 RACK1-interacting proteins shared within all the four infection time points (Fig. 1A). Further gene ontology (http://amigo.geneontology.org/amigo/landing) pathway analysis (Fig. 1B) of the 179 overlapping proteins showed that integrin signaling pathway (including ITGB3 and its related factors such as ILK, ACTN1, PIK3C2A, MAPK3 and FYN) is one of the most enriched pathways. As strong evidence has demonstrated that RACK1 could interact with a number of integrins during various physiological processes and pathogenesis (19, 24, 31, 38), ITGB3 was therefore chosen for the following studies. The mRNA and amino acids sequences of ITGB3 from eighteen species were downloaded for phylogenetic analysis using online tool Clustal Omega (https://www.ebi.ac.uk/Tools/msa/clustalo/) with the default parameters (Supplementary Fig. 1A and 1B). The ITGB3 amino acid sequence is highly conserved between human, monkey and pig (Supplementary Fig. 2).

**Figure 1.**
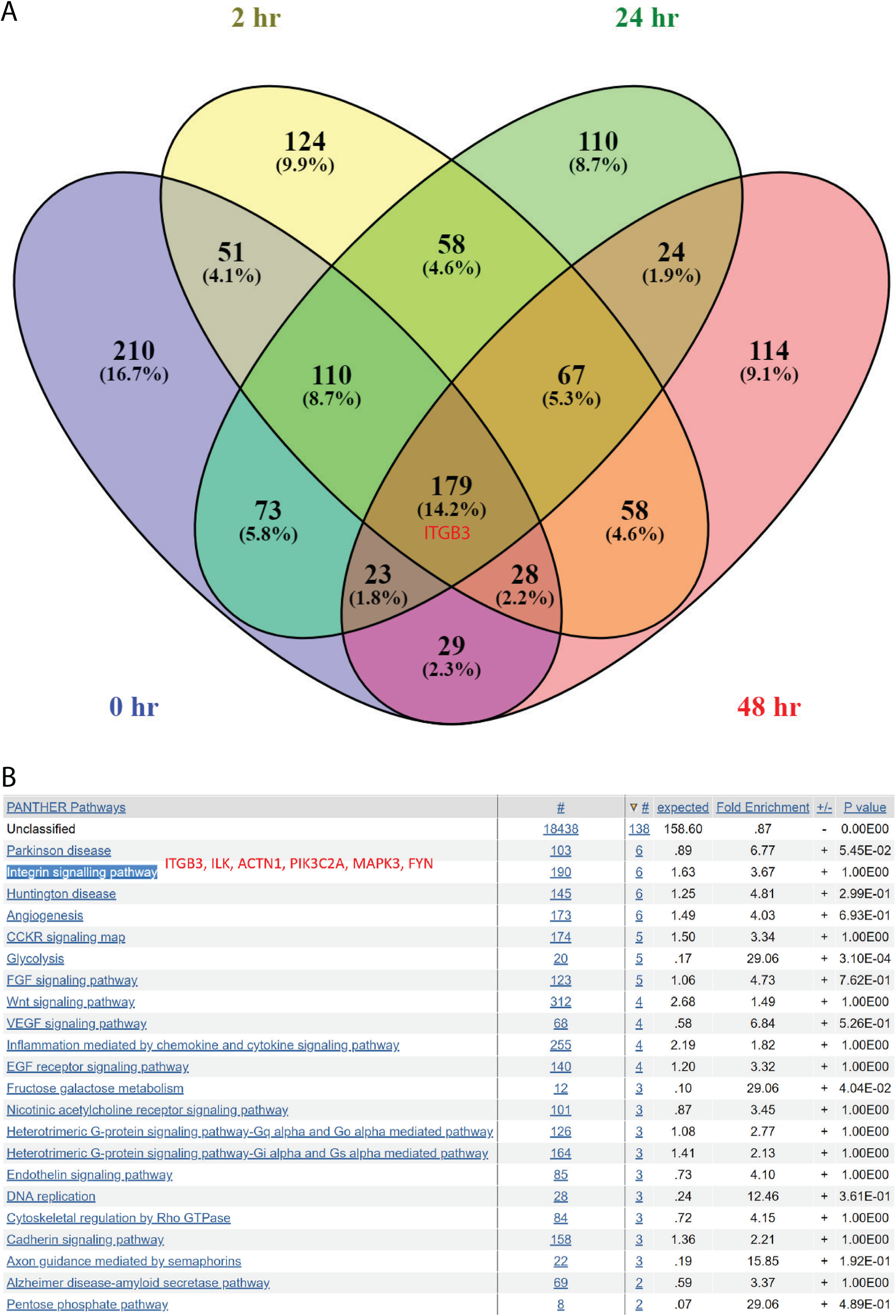
Identification of ITGB3 as a RACK1 interacting protein. A. After pulldown of overexpressed RACK1 from Marc-145 cells infected with PRRSV YN-1 strain, potential RACK1 interacting co-precipitants were subjected to mass spectrometry identification. Venny diagram indicates the overlapping among the samples from different time points (0 hr: Blue, 2 hr: Yellow, 24 hr: Green and 48 hr: Red) post infection. Totally 179 proteins, including ITGB3 (indicated in red), were shared by all four samples. B. Gene ontology (http://amigo.geneontology.org/amigo/landing) analysis of the 179 common proteins revealed that integrin signaling pathway (comprising of ITGB3, ILK, ACTN1, PIK3C2A, MAPK3 and FYN, highlighted in red) was one of the top enriched pathways.

### Abrogation of ITGB3 inhibited PRRSV replication and NF-κB activation in Marc-145 cells

Infection of Marc-145 cells with PRRSV YN-1 strain (100 TCID_50_) activated NF-κB signal pathway, as reflected by phosphorylation of IκBα (Fig. 2G) and p65 (Fig. 2E), in line with our previous studies (26, 28). Compared with the non-infection group, a striking upregulation of ITGB3 (Fig. 2C) was observed during the whole experimental time period (from 1 to 60 hours post inoculation), even at the very beginning of PRRSV infection (such as 1–12 hpi).

**Figure 2.**
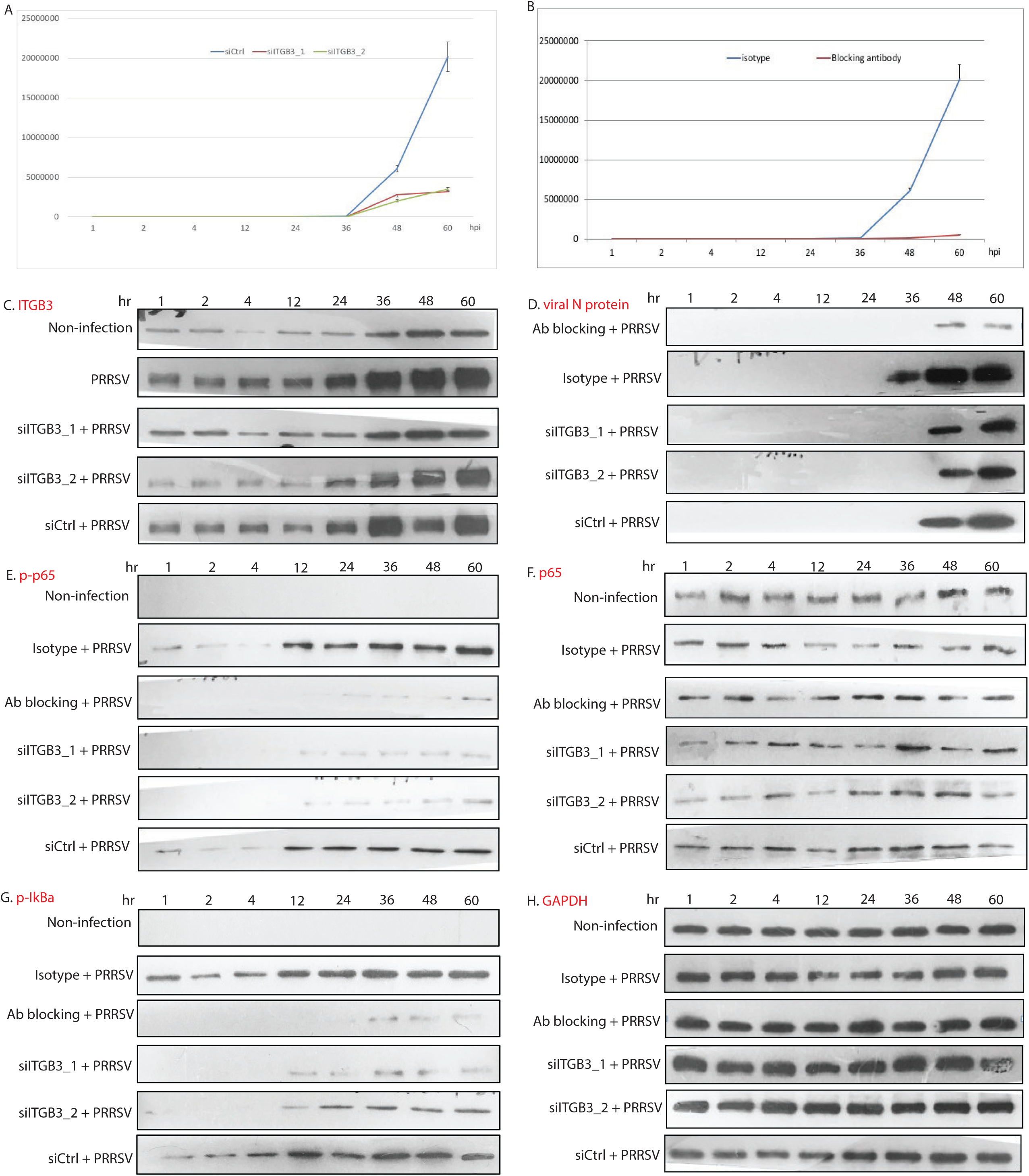
Abrogation of ITGB3 inhibited PRRSV infection and NF-κB activation in Marc-145 cells. mRNA expression level of viral ORF7 was measured by RT-qPCR and normalized to cellular GAPDH after different time points post abrogation of ITGB3 through siRNA knockdown (A) or antibody blocking (B). Forty-eight hours post siRNA knockdown or two hours post antibody blocking, Marc-145 cells were infected with PRRSV YN-1 strain (100 TCID_50_). Total protein samples were collected at the indicated infection time points (1, 2, 4, 12, 24, 36, 48 and 60 hpi) and analyzed by western blot for ITGB3 (C), viral N protein (D), phosphorylated-p65 (E), p65 (F), phosphorylated-IκBα (G) and GAPDH (H).

Abrogation of ITGB3 by siRNA knockdown or antibody blocking markedly abolished the PRRSV infection in Marc-145 cells, reflected on both viral mRNA (Fig. 2A and 2B) and protein (Fig. 2D) levels. Meanwhile, inhibition of phosphorylation of IκBα (Fig. 2G) and p65 (Fig. 2E) by annulment of ITGB3 suggested obliteration of NF-κB activation upon PRRSV infection, while not affecting the protein level of p65 (Fig. 2F) and GAPDH (Fig. 2H). Moreover, antibody blocking showed more pronounced inhibitory effects on PRRSV replication and NF-κB induction.

### Overexpression of ITGB3 enhanced PRRSV replication and NF-κB activation in Marc-145 cells

Transfection of ITGB3 expression vector increased the ITGB3 level, while combination with PRRSV further elevated the ITGB3 expression (Fig. 3A). Overexpression of ITGB3 also induced the phosphorylation of IκBα (Fig. 3B) and expression of RACK1 (Fig. 3C), but not p65 (Fig. 3D). Compared with the corresponding non-infection controls, further elevations of p-IκBα (Fig. 3B), RACK1 (Fig. 3C) and p-p65 (Fig. 3E) were observed, but the expression level of p65 (Fig. 3D) or GAPDH (Fig. 3F) was not enhanced. Viral N protein was detectable at 48 hours post infection (hpi), with a weak band was observable already 36 hpi (Fig. 3G) when ITGB3 was overexpressed, indicating a boosted PRRSV infection by ITGB3 upregulation, which is confirmed by the immunoflueorescence data (Fig. 4).

**Figure 3.**
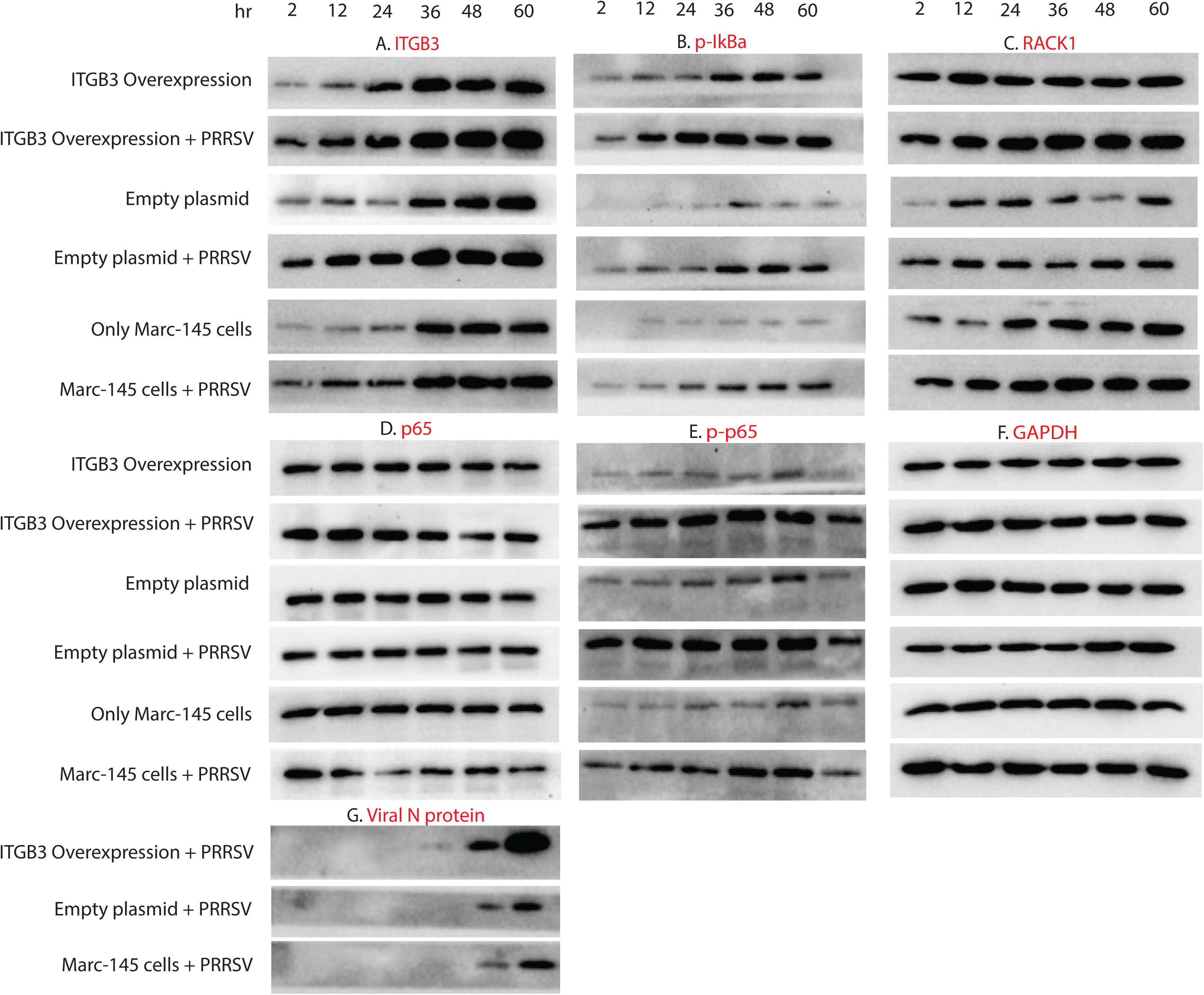
Overexpression of ITGB3 enhanced PRRSV replication and NF-κB activation in Marc-145 cells. Twenty-four hours post plasmid transfection to overexpress ITGB3, Marc-145 cells were infected with PRRSV YN-1 strain (100 TCID_50_). Total protein samples were collected at the indicated infection time points (2, 12, 24, 36, 48 and 60 hpi) and analyzed by western blot for ITGB3 (A), phosphorylated-IκBα (B), RACK1 (C), p65 (D), phosphorylated-p65 (E), GAPDH (F) and viral N protein (G).

**Figure 4.**
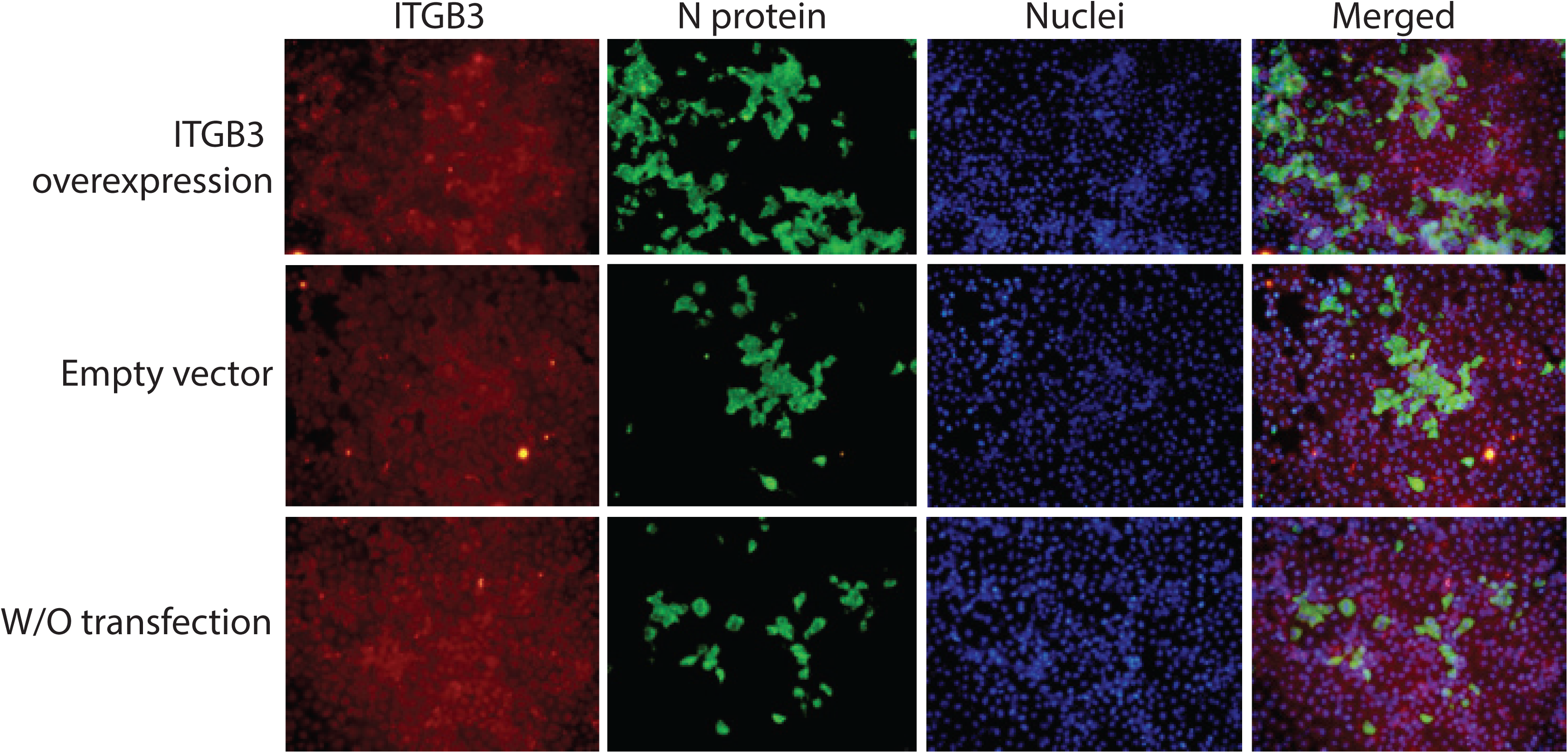
Indirect immunofluorescence detection of enhanced PRRSV replication in Marc-145 cells with ITGB3 overexpression. Transfection of the ITGB3 overexpression plasmid in 6-well plate was performed 24 hours prior to infection with PRRSV YN-1 strain (2000 TCID50/well). Cells were fixed 48 hours post infection. Anti-N monoclonal antibody (mAb) and Alexa Fluo488 conjugated secondary antibody were applied in indirect immunofluorescence staining to demonstrate the viral N protein level, with anti-ITGB3 polyclonal antibody and Alexa Fluo546 conjugated secondary antibody to visualize the ITGB3 protein level. Nuclei are shown in blue, ITGB3 protein in red, and viral N protein in green. The images of the same treatment from the three channels were merged. These images are representative of three independent experiments.

### Loss-of-function of ITGB3 protected Marc-145 cells from cytopathic effects and reduced the viral titer

Change of cytopathic effects (CPE) and viral titer are among the most direct and undoubted parameters in virus research. We found that compared with the controls (non-treatment, siCtrl and Isotype antibody) where severe CPEs were manifested till dilution of 10^−5^, loss-of-function of ITGB3 by siITGB3 or antibody blocking could significantly protect Marc-145 cells from CPE at the viral dilution between 10^−3^ and 10^−4^ (Fig. 5). The TCID_50_ titer of PRRSV was reduced by 40-72 times when ITGB3 was knocked-down or by approximate 300 times when blocked with ITGB3 specific antibody (Fig. 5).

**Figure 5.**
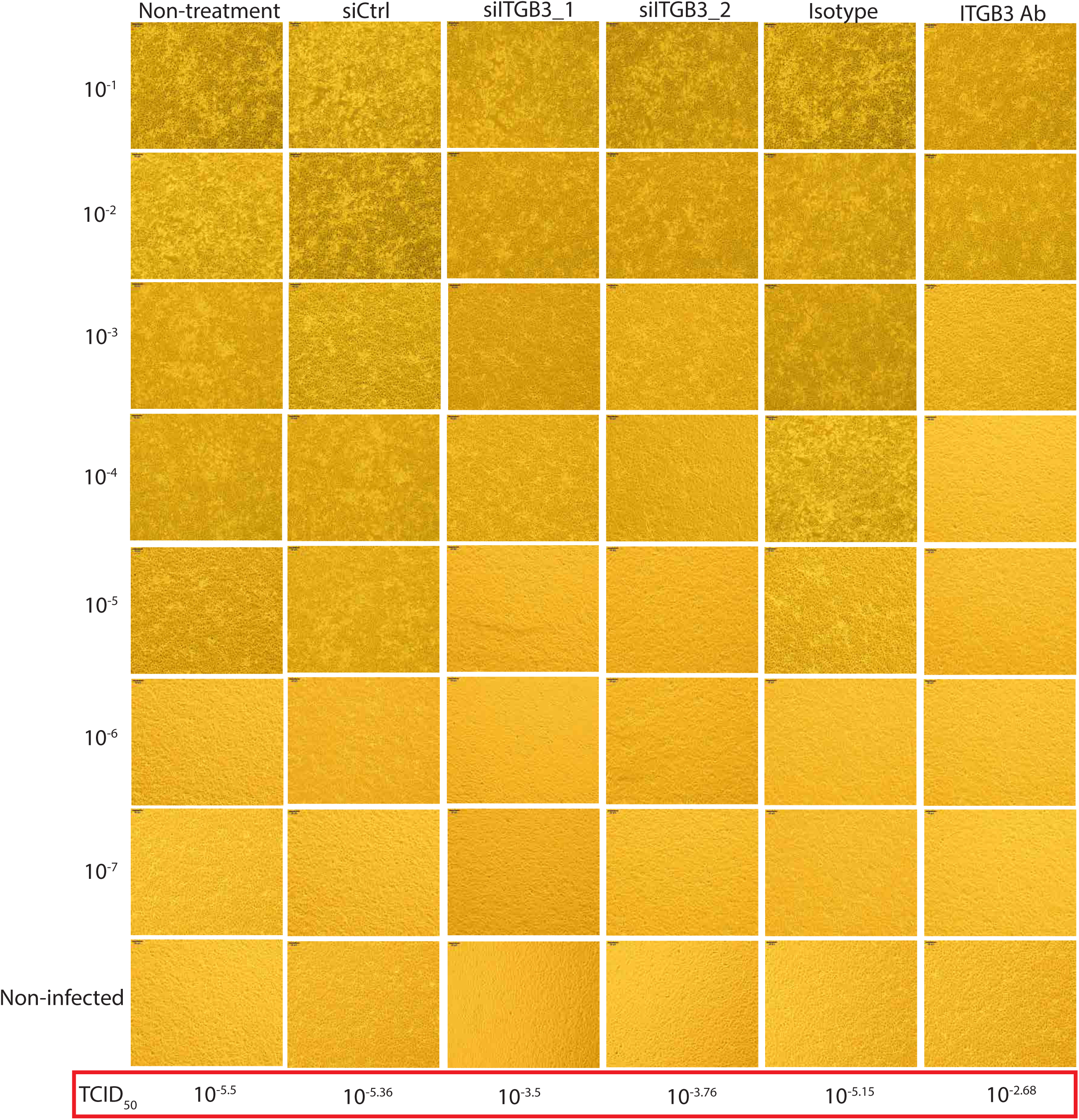
The siRNA knockdown or antibody blocking of ITGB3 shielded Marc-145 cells from cytopathic effects (CPE). CPE was recorded using the inverted microscope over a period of 3 days post infection. The 50% tissue culture infected dose (TCID_50_) was determined by Reed–Muench method. Compared with the corresponding control group, the TCID_50_ viral titer was elevated by 40-70 times (siRNA knockdown) or approximately 300 times (blocking with ITGB3 specific antibody) respectively (highlighted in red square). This data is representative for three independent experiments. The bar in each microscopic image (100 × magnification) indicates 50 μm.

### RACK1 and ITGB3 are NF-κB target genes

The mouse RACK1 gene was reported to be regulated by NF-κB (39), however, there is no evidence to show whether the RACK1 and ITGB3 from monkey kidney Marc-145 cell line are target genes of NF-κB. As expected, besides the phosphorylation of IκBα and p65 (Fig. 6E and 6G), which confirmed our previous findings (26, 28), PRRSV infection enhanced the expression of ITGB3 and RACK1 (Fig. 6A and 6B). Additionally, phosphorylation of TRAF2 (Fig. 6D) was upregulated after PRRSV infection, while without noticeably changing the expression of TRAF2 (Fig. 6C) and p65 (Fig. 6F). When NF-κB signaling pathway inhibitor BAY 11-7082 was introduced, the expression of RACK1 (Fig. 6A) and ITGB3 (Fig. 6B), with or without PRRSV infection, were suppressed, indicating that RACK1 and ITGB3 are the target genes of NF-κB. When NF-κB was inhibited, the PRRSV N protein was barely detectable at 48 hpi, compared with a clear band in the un-treated group (Fig. 6H), implying an important role of NF-κB activation in PRRSV infection.

**Figure 6.**
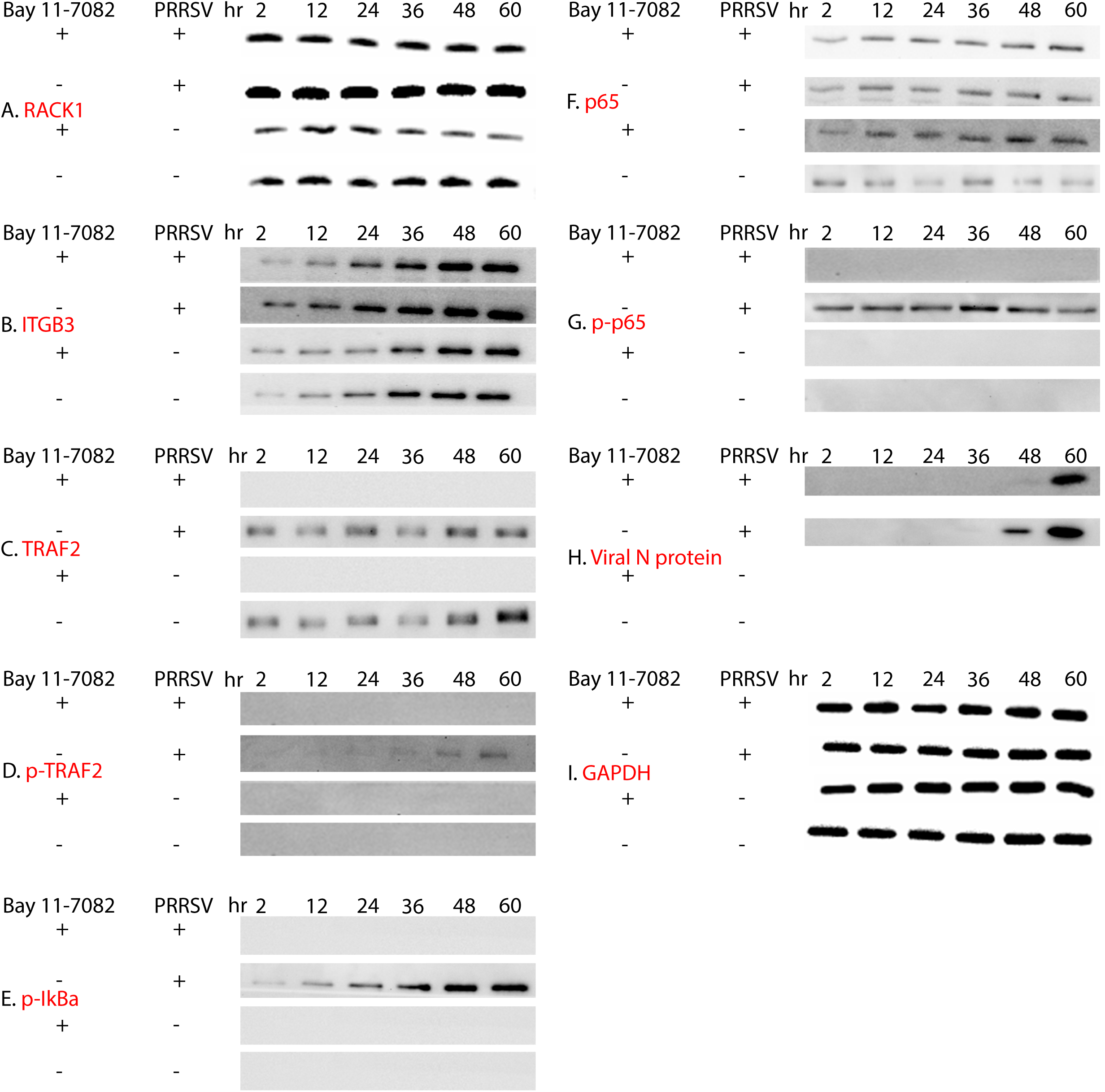
RACK1 and ITGB3 are NF-κB target genes and treatment with NF-κB inhibitor BAY 11-7082 alleviates PRRSV replication in Marc-145 cells through TRAF2 and its phosphorylation. Marc-145 cells were infected with PRRSV YN-1 strain two hours after or without pre-treatment with BAY 11-7082. The cells at different time points (2, 12, 24, 36, 48 and 60 hpi) were collected for western blot against RACK1 (A), ITGB3 (B), TRAF2 (C), phosphor-TRAF2 (D), phosphorylated-IκBα (E), p65 (F), phosphorylated-p65 (G), viral N protein (H) and GAPDH (I). Non-infected cells were applied as control.

### ITGB3 and RACK1 regulated each other

Compared with the neutral control (siCtrl), significant knockdown was achieved through specific siRNAs for both ITGB3 (Fig. 7A) and RACK1 (Fig. 7B) over the experiment time period (from 1 to 60 hpi). Furthermore, knockdown of RACK1 resulted in faint bands of ITGB3 (Fig. 7A), meanwhile, weak bands of RACK1 were also observed upon knockdown of ITGB3 (Fig. 7B), although the phenotype was not as prominent as from knockdown of the corresponding gene itself. In addition, ITGB3 knockdown diminished the upregulation of ITGB3 by PRRSV infection, and that of RACK1 to a much less extent. Likewise, inhibition of RACK1 lessened the induction of both RACK1 and ITGB3 by PRRSV infection. Taken together, the data suggested the inter-regulation of ITGB3 and RACK1.

**Figure 7.**
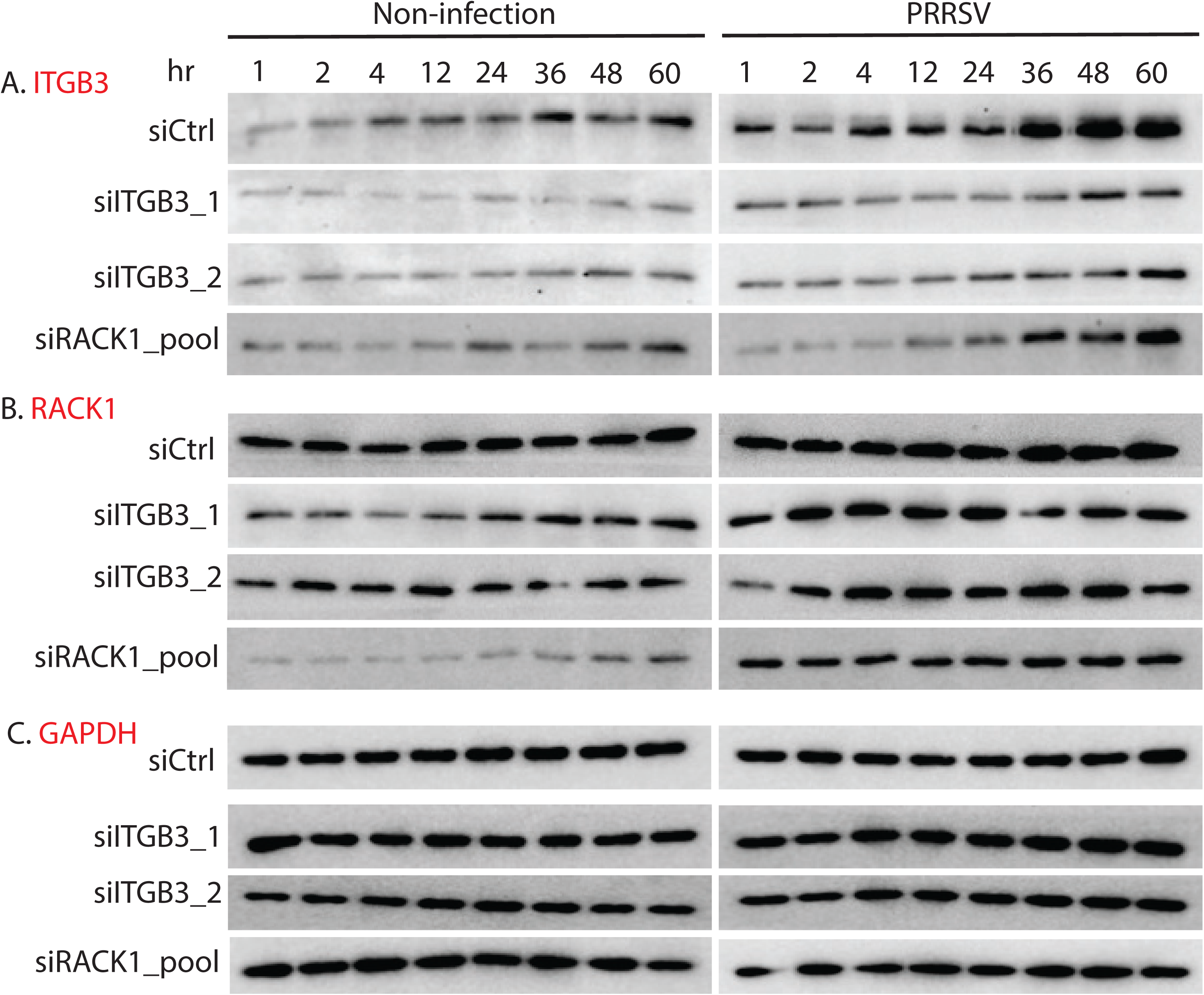
ITGB3 and RACK1 regulated each other. Forty-eight hours post siRNA knockdown (non-targeting control, siITGB3 and siRACK1), Marc-145 cells were infected with PRRSV YN-1 strain (100 TCID_50_) or without infection. Total protein was collected at the indicated infection time points (1, 2, 4, 12, 24, 36, 48 and 60 hpi) and analyzed by western blot for ITGB3 (A), RACK1 (B) and GAPDH (C).

## DISCUSSION

RACK1 is well known to interact with the membrane proximal region of the cytoplasmic tail of integrins β1 (20, 24), β2 (40), β3 (38) and β5 (31). The first identification of integrin β3 (ITGB3) as a RACK1-interacting protein by pulldown and mass spectrometry in Marc-145 cell in this study is consistent with previous discoveries in other species. Besides, some other RACK1 interacting and ITGB3 related proteins (e.g., ILK, ACTN1, PIK3C2A, FYN and ERK1) are enriched in the integrin signaling pathway and worth further investigation.

The primary target cells of PRRSV infection *in vivo* are PAMs. However, due to lack of stable and immortal PAM cell lines, Marc-145 cells are extensively used for PRRSV basic research as it is highly permissive for PRRSV infection. In this study, PRRSV infection in Marc-145 cells led to the upregulation of the ITGB3 expression. Overexpression of ITGB3 in turn elevated PRRSV replication and strengthened the NF-κB activation. Abolishment of ITGB3 repressed PRRSV replication and inhibited the NF-κB activation. Furthermore, downregulation of ITGB3 protected the Marc-145 cells from the cytopathic effects and abridged the viral titer. All these results indicated the pivotal role of ITGB3 in PRRSV infection, with the underlying mechanisms for further investigations.

Vimentin was identified as an important part of the PRRSV receptor complex in Marc-145 (41). It has been reported that the vimentin head domain bound to the ITGB3 cytoplasmic tail (42). As a cell surface protein, ITGB3 is known to be involved in the cell entry of West Nile Virus (43), Herpes Simplex Virus (44), Japanese Encephalitis Virus (45) and Foot-and-Mouth Disease Virus (46). It is logical to hypothesize that ITGB3 is also a part of the PRRSV receptor complex and play its roles in the attachment and entry of PRRSV. More interestingly, the infection of PRRSV displayed positive feedback effect on the expression of ITGB3, as observed also in Swine Fever Virus (47, 48) and Dengue Virus (49).

Activation of NF-κB is a hallmark of most viral infections and the viral pathogens in many cases hijack NF-κB pathway to their own advantages (50). Suppression of NF-κB and the subsequent delay of detectable viral mRNA and N protein after ITGB3 knockdown implied that the NF-κB activation after PRRSV is beneficial for its infection/replication. Previous study showed that the mouse *RACK1* gene was regulated by NF-κB activated by nerve growth factor (NGF) in PC-12 cells (39). In line with that, we showed that monkey *RACK1* was a NF-κB target gene during PRRSV infection in Marc-145 cells. Moreover, our data for the first time indicated that ITGB3 was a novel NF-κB target gene. Presumably, PRRSV infection in Marc-145 cells activated NF-κB signal pathway to transcribe its target genes, including RACK1 and ITGB3, which in turn facilitated the PRRSV infection/replication.

As a NF-κB inhibitor, BAY 11-7082 has been widely used to investigate the effects of NF-κB inhibition on diverse physiological and pathogenic processes (51), as well as in PRRSV infection (52). However, inhibition of NF-κB signal pathway alone by BAY 11-7082 could not completely nullify the PRRSV replication in Marc-145 cells (Fig. 6H). This suggests that the parameters for NF-κB inhibition (concentration and time) were not optimal or/and that NF-κB activation might only partially contributes to PRRSV infection. We speculate that some other signal pathways may be involved in and indispensable for PRRSV infection. Indeed, there have been a number of studies illustrating the relationship between PRRSV infection and mTOR signaling pathway (53), MyD88/ERK/AP-1 and NLRP3 inflammasome (54), ERK1/2-p-C/EBP-β Pathway (55) and Wnt/β-catenin pathway (56). Simultaneously deactivating the above-mentioned signal pathways probably will further block the PRRSV infection.

Similarly, NF-κB inhibition could not completely block the expression of ITGB3 or RACK1 induced by PRRSV infection, implying the existence of bypass signal pathways activated by PRRSV infection. Another possibility could be that within the NF-κB pathway, RACK1 and ITGB3 do not directly regulate each other, but via upstream/downstream pattern. To further complicate the situations, there can be some other regulatory molecules mediating RACK1 and ITGB3. As a receptor for activated protein kinase C, RACK1 was shown to form a phorbol 12-myristate 13-acetate (PMA)-induced complex between PKCε and integrin β1 or β5 in human glioma cells (38). Human articular chondrocytes express PKCα, γ, δ, ι and λ, and upon stimulation chondrocytes show a rapid, integrin β1 dependent, translocation of PKCα to the cell membrane and increased association of RACK1 with PKCα and β1 integrin (57). We therefore contemplate one or more PKC members may play some certain roles together with RACK1 and ITGB3 during the NF-κB pathway induced by PRRSV infection.

In conclusion, ITGB3 plays an imperative role during PRRSV infection and NF-κB activation in Marc-145 cells, presumably as a co-receptor for example with vimentin or CD163; PRRSV infection activates a positive feedback loop involving the activation of NF-κB and upregulation of ITGB3 and RACK1 in Marc-145 cells (Fig. 8). However, the mechanistic linkage among PRRSV infection, the expression and interaction between RACK1 and ITGB3, and the NF-κB signaling pathway desires further investigations. Our data would shed new light on the underlying mechanisms of PRRSV infection in swine and suggested RACK1-ITGB3-NF-κB signaling axis as a promising potential therapeutic target for PRRS control.

**Figure 8.**
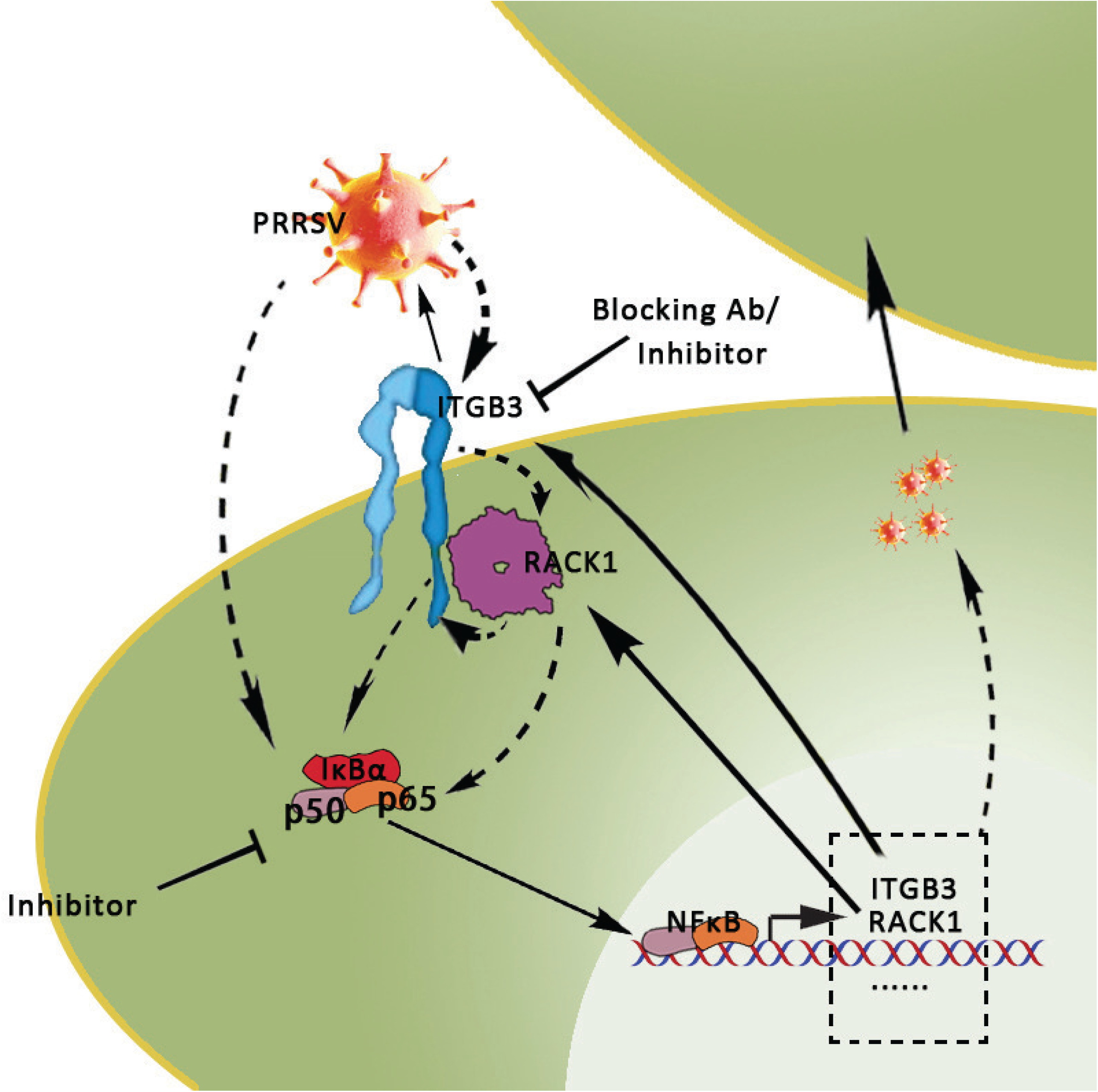
Schematic diagram of presumed host–pathogen interaction mechanisms. ITGB3 interacts with RACK1 and facilitates PRRSV infection and NF-κB activation in Marc-145 cells, presumably as a co-receptor of CD136 or vimentin. PRRSV infection activates a positive feedback loop involving activation of NF-κB and upregulation of ITGB3 and RACK1. Both ITGB3 and RACK1 were NF-κB target genes during PRRSV infection, and they regulate each other. The activation of NF-κB and transcription of its downstream genes are beneficial for PRRSV infection/replication. ITGB3 and NF-κB signaling pathway are potential therapeutic targets for PRRS control.

## MATERIAL AND METHODS

Most of the materials and methods used in this study were described with details in our previous studies (26, 28, 58, 59).

### Virus and cells

The highly pathogenic PRRSV strain YN-1, which belongs to the PRRSV genotype 2 (the North American genotype), was isolated by our lab (TCID_50_= 10^−5.5^, GenBank accession number KJ747052). The highly PRRSV permissive grivet monkey kidney derived Marc-145 cells were used as cell model in this study.

### Pulldown and mass spectrometry analyses of RACK1 binding partners

Briefly, full length *RACK1* cDNA was cloned from Marc-145 cell with the primer pair RACK1-F and RACK1-R (listed in Table 1). After sequencing confirmation, the cDNA was subcloned into a prokaryotic expression vector (pET32a, EMD Biosciences) with a 6xHis tag. Recombinant His-RACK1 bait protein was expressed in *E.coli* BL21 (DE3) cells and incubated with Ni-NTA resin. Meanwhile, Marc-145 cells were infected with 100 TCID_50_ for 2 hr, 24 hr and 48 hr, respectively, with non-infected cells (0 hr) as control. Sufficient amount of total protein (100 µg/ml) from each group was extracted and loaded to the affinity column containing His-RACK1 which were bound to Ni-NTA for affinity Pulldown of target proteins. After washing, the isolated proteins were eluted and identified by mass spectrometry. More detailed protocol is included in the Supplementary Material and Methods.

**Table 1.**
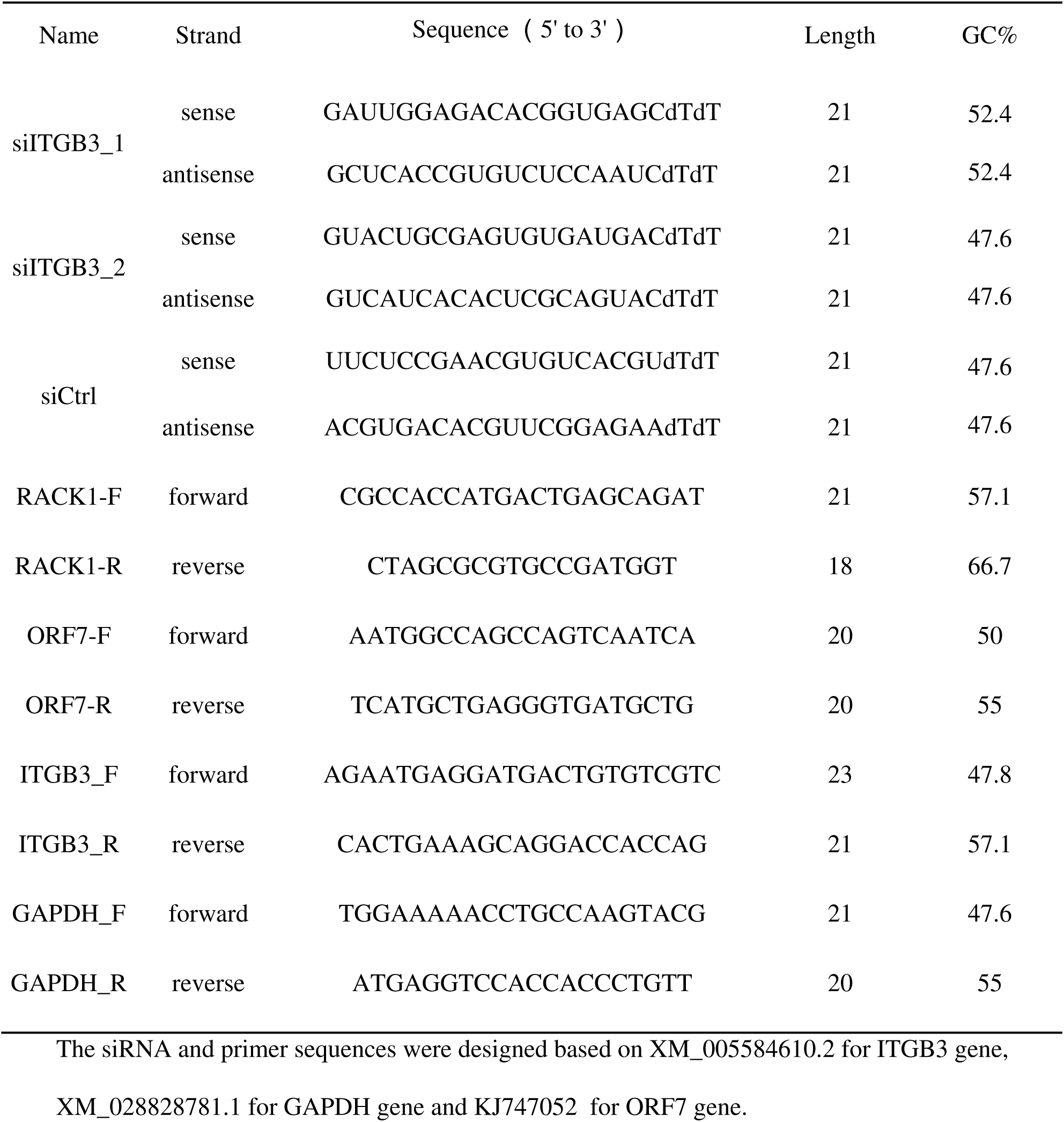
siRNA and primer sequences used in this study

### siRNA transfection

Transfections were performed in 6-well plates with 3×10^5^ cells seeded into 2 ml complete medium forty-eight hours before infection with YN-1 strain. Forty nM siRNAs against ITGB3 or RACK1 or non-targeting siRNA (sequences listed in Table 1, purchased from Sangon, Shanghai, China) were premixed with 6 µl Lipofectamine 3000 in OptiMEM and added to each well. siRACK1-pool is the mixture of the two siRNAs with equal concentration against RACK1 used in our previous study (26). Transfection with non-targeting siRNA (siCtrl) was used as neutral control for normalization or comparison with the specific knockdowns.

### Treatment with ITGB3 antibody and NF-κB inhibitor BAY 11-7082

Briefly, 3×10^5^ Marc-145 cells in 2 ml cell culture medium were seeded into 6-well plate and cultured overnight. Then the cells were incubated with either an antibody specifically binding to ITGβ3 or a Mouse mAb IgG2b Isotype control (listed in Table 2) at 25 μg/ml (45) for 2 h at 37 °C, followed by medium change and PRRSV infection. For NF-κB inhibition, Marc-145 cells were pre-treated with 10 μM BAY 11-7082 (#B5556-10MG, Sigma-Aldrich) for 2 hours, followed by medium change and PRRSV infection. DMSO at 10 μM was applied as negative control.

**Table 2.**
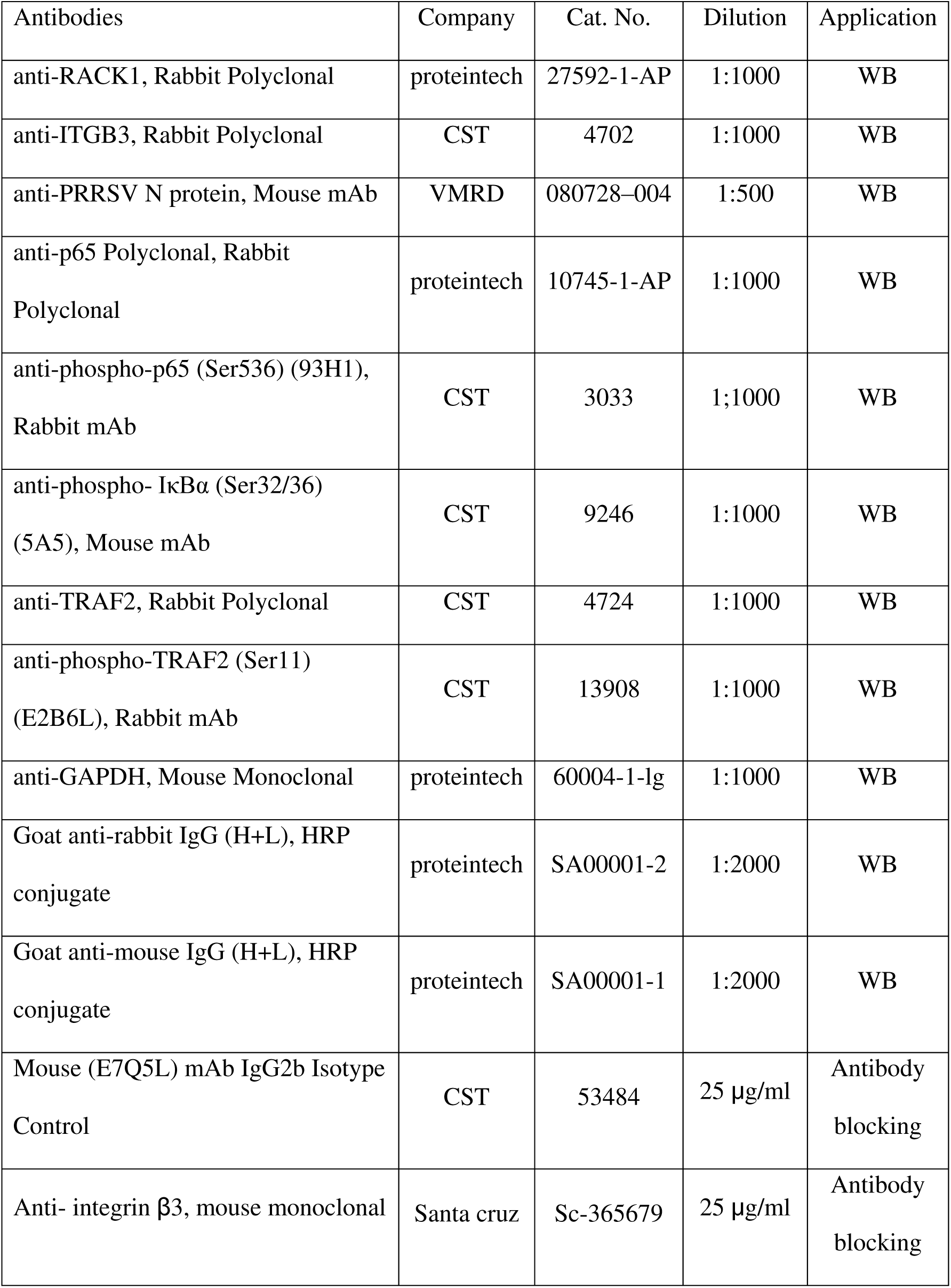
Antibodies used in this study

| Antibodies | Company | Cat. No. | Dilution | Application |
| --- | --- | --- | --- | --- |
| anti-RACK1, Rabbit Polyclonal | proteintech | 27592-1-AP | 1:1000 | WB |
| anti-ITGB3, Rabbit Polyclonal | CST | 4702 | 1:1000 | WB |
| anti-PRRSV N protein, Mouse mAb | VMRD | 080728-004 | 1:500 | WB |
| anti-p65 Polyclonal, Rabbit Polyclonal | proteintech | 10745-1-AP | 1:1000 | WB |
| anti-phospho-p65 (Ser536) (93H1), Rabbit mAb | CST | 3033 | 1;1000 | WB |
| anti-phospho- IκBα (Ser32/36) (5A5), Mouse mAb | CST | 9246 | 1:1000 | WB |
| anti-TRAF2, Rabbit Polyclonal | CST | 4724 | 1:1000 | WB |
| anti-phospho-TRAF2 (Ser11) (E2B6L), Rabbit mAb | CST | 13908 | 1:1000 | WB |
| anti-GAPDH, Mouse Monoclonal | proteintech | 60004-1-Ig | 1:1000 | WB |
| Goat anti-rabbit IgG (H+L), HRP conjugate | proteintech | SA00001-2 | 1:2000 | WB |
| Goat anti-mouse IgG (H+L), HRP conjugate | proteintech | SA00001-1 | 1:2000 | WB |
| Mouse (E7Q5L) mAb IgG2b Isotype Control | CST | 53484 | 25 μg/ml | Antibody blocking |
| Anti- integrin β3, mouse monoclonal | Santa cruz | Sc-365679 | 25 μg/ml | Antibody blocking |

### Virus infection

Forty-right hours post transfection with siRNA or overexpression plasmids, or two hours post antibody blocking or NF-κB inhibition, the Marc-145 cells were inoculated with 100 TCID_50_ PRRSV YN-1 strain per well for 2 hrs. Then the virus was withdrawn by medium change and the cells were kept in culture till further analysis.

### Total RNA isolation and RT-qPCR analysis

Total RNA was isolated using RNAiso Plus RNA isolation kit (#9108/9109, Takara, Dalian, China) from Marc-145 cells at different time points (1, 2, 4, 12, 24, 36, 48 and 60 hpi) according to the manufacturer’s instructions. RT-qPCR was performed to determine the relative expression level of viral ORF7 gene for quantification with cellular GAPDH served as an internal control (primer sequences are listed in Table 1).

### Overexpression of ITGB3 in Marc-145 cells

Briefly, *ITGB3* cRNA was reverse transcribed and amplified from Marc-145 total RNA with three pairs of primers (listed in Supplementary Materials and Methods) designed according to its sequence (GenBank seq. No. XM_005584610.2) and resulted in three fragmented sequences with restrict enzyme adapter at each end. After sequencing confirmation, the ITGB3 fragments were enzymatically digested, purified and ligated to pcDNA™3.1/Hygro(+) mammalian expression vector (#V87520, Thermofisher Scientific, MA, USA). More detailed protocol is included in the Supplementary Materials and Methods.

Marc-145 cells (3×10^5^) were cultured in 2 ml medium in 6-well plates until 70-80% confluence, followed by transfection with 3 μg expression plasmid (pcDNA3.1-ITGB3) or empty vector (pcDNA3.1) and 3.75 µl Lipofectamine 3000. Twenty-four hours post transfection, the cells were inoculated with PRRSV YN-1 strain (2000 TCID_50_/well, equivalent to an approximate MOI of 5) till further analysis. Transfection with empty vector and cells without infection were used as controls for normalization or comparison with the specific overexpression of ITGB3.

### Western blot analysis

Total viral and cellular proteins were extracted with RIPA buffer at the indicated time points and denatured immediately by boiling in SDS-PAGE loading buffer. After separated by electroporation and transferred to nitrocellulose membrane, the proteins of interest were detected with antibodies as listed in Table 2 at indicated dilutions. The blots were developed with chemiluminescent ECL Plus substrate and visualized using chemiluminescent film.

### Indirect immunofluorescence staining

Twenty-four hours post transfection of ITGB3 overexpression plasmid, the cells were inoculated with PRRSV YN-1 strain (2000 TCID_50_/well). Transfection with empty vector and cells without infection were used for comparison. Forty-eight hours post PRRSV infection, the Marc-145 cells were washed with PBS (0.05 M, pH7.4) and fixed with 4% paraformaldehyde (PFA) at room temperature for 10–15 minutes. Then the cells were washed with PBS for three times, permeabilizated with PBS containing 0.3% Triton X-100 for 15 minutes and blocked with PBS containing 1% BSA for 2 hours at 4 °C. The nuclei staining with 5 μg/ml of Hoechst 33342 (Sigma-Aldrich, Cat. #B2261) was carried out for 20 minutes at room temperature to visualize the nuclei. The cells were subsequently co-incubated with PRRSV antibody against N protein and anti-ITGB3 antibody at 4 °C overnight, washed three times with PBS and then co-incubated with corresponding secondary antibody for 1 h at 37 °C (all antibody information is listed in Table 2). After three times PBS wash, the cells were subjected to image analysis by fluorescence microscopy (Olympus).

### TCID_50_ measurement after ITGB3 siRNA transfection and antibody blocking

Marc-145 cells were seeded into 96-well plates with10000 cells per well. Forty-eight hours post siRNA knockdown or 2 hours post antibody blocking, a 10× serial dilution of PRRSV YN-1 strain was prepared and added into the cells with hexatruple replicates in each concentration. Cytopathological effects (CPE) were monitored using the inverted microscope over 3 days post infection. Well number was counted and the 50% tissue culture infected does (TCID_50_) was determined by Reed-Muench method. Non-treated cells, cells transfected with non-targeting siRNA or blocked with unspecific IgG antibody were used as control.

## Declaration of interest

None.

## Acknowledgements

This work was supported by the National Natural Science Foundation of China (grant no. 31560705 and 31960701), by the Key Projects of Yunnan Provincial Natural Science Foundation (No. 2016FA018), by Program for Innovative Research Team (in Science and Technology) in University of Yunnan Province (IRTSTYN) and by Pig Disease Research Center, Yunnan Agricultural University. The funders had no role in study design, data collection and interpretation, or the decision to submit the work for publication.

## Author contributions

Chao Yang and Rui Lan performed all the western blots related experiments and overexpression of ITGB3. Xiaochun Wang and Qian Zhao performed the overexpression of RACK1 and subsequent pulldown. Xidan Li and Junlong Bi performed all the data analysis. Jing Wang and Guishu Yang performed the RT-qPCR and immunofluorescence staining. Yingbo Lin generated the mechanistic hypothesis and modified the manuscript. Jianping Liu and Gefen Yin supervised this study and wrote the manuscript.

## Appendix A. Supplementary data

Supplementary data related to this article can be found in the online version.

---

Supplementary Figure 1. Phylogenetic analysis of ITGB3 mRNA (A) and amino acid (B) sequences from eighteen species. The sequence clustering was performed using the online tool Clustal Omega (https://www.ebi.ac.uk/Tools/msa/clustalo/). Three species including pig, human and Rhesus monkey are highlighted in red.

Supplementary Figure 2. Multiple sequence alignment of ITGB3 amino acid sequences from pig, human and Rhesus monkey (corresponding to Marc-145 cells, derived from African green monkey kidney) showed high similarity. Positions sharing a single, fully conserved residue are indicated with an asterisk (*), while conservations between groups of strongly similar properties with scoring > 0.5 and =< 0.5 in the Gonnet PAM 250 matrix are indicated with colon (:) and dot (.), respectively.

